## Supplementary Information for "The evolution of trait correlations constrains phenotypic adaptation to high CO_2_ in a eukaryotic alga"

Nathan G. Walworth[1], Jana Hinners[2], Phoebe Argyle[3], Suzana G. Leles[1], Martina A. Doblin[3], Sinéad Collins[2], Naomi M. Levine[1]*

[1]Department of Biological Sciences, University of Southern California, Los Angeles, California 90089-0371, USA, [2] Institute of Evolutionary Biology, University of Edinburgh, Edinburgh EH9 3FL, United Kingdom, [3] Climate Change Cluster, University of Technology Sydney, Sydney, NSW 2007, Australia

**Corresponding author**

**Supplementary Information 1:** ***Adaptation and population size***

Adaptation dynamics in TRACE is based on an adapted Fisher's geometric adaptation model from Kronholm and Collins [1] and Walworth et al 2020 [2]. The model simulates a population that initially starts far from a high-fitness optimum. Adaptation progresses in the model through predominantly non-neutral changes to traits and/or correlations in the population which generates an adaptive walk towards the high-fitness optimum. We do not explicitly represent drift in the model, which can also impact the adaptive process. However, the relative supply of changes (population size x number of changes per generation) used in our model runs is sufficient for selection to overwhelm drift resulting in robust evolutionary results. Accordingly, in prior studies using an analogous adaptive process, we varied population size and/or selection strength over multiple orders of magnitude and demonstrated adaptive outcomes to be robust over the same selection period [1,2].

**Supplementary Information 2:** ***Sensitivity tests***

We conducted several sensitivity tests to examine model dynamics. First, we tested if the sequence of correlational changes influenced adaptive outcomes in our model. To do this, we conducted model runs where we changed the order in which traits were updated and showed that phenotypic results remained unchanged as expected (Supplementary Fig. 8).

We also examined the impact of trait correlational constraints on evolutionary trajectories by randomly changing traits independently of trait correlations (i.e. ignoring trait relationships) (Supplementary Fig. 2). For these runs, each individual experienced a random trait change as in the default model but no other traits were updated. Hence, individuals were unconstrained by bias and so were able to quickly move directly to the high-fitness area. We found that only one phenotype emerged, as expected (Supplementary Fig. 2). This demonstrates that trait correlational constraints are critical for producing different evolutionary strategies (i.e. emergent, cryptic phenotypes), and if constraints are not present, individuals are able to more freely explore phenotypic space and arrive at the high-fitness phenotype more rapidly.

Finally, we tested the sensitivity of the model dynamics by varying the ratios of trait and trait correlation changes (Supplementary Fig. 9). In addition to the default Mode 90/10 (90% of individuals experience a trait change only and 10% experience both a trait + trait correlation change) two other modes were run: Mode 50/50 and Mode 10/90. For each generation in Mode 50/50, 50% of individuals experienced a trait change and 50% experienced both a trait and correlation change. For Mode 10/90, 10% of individuals experienced a trait change and 90% of individuals experienced both a trait and correlation change. All other parameters stayed the same. Mode 50/50 found the same 4 phenotypes as the default Mode 90/10 while Mode 10/90 found the two most accessible phenotypes, Pop-MA and Pop-MD (Supplementary Fig. 9). No new populations emerged. The fact that some combination of the same phenotypes emerged from each of the three independently run modes provides further evidence for a robust, conserved trait-scape with limited high-fitness phenotypes derived from a population with no historical bias.

**Supplementary Information 3:** ***Supplemental Methods***

For all runs, we used the same evolved trait-scape (i.e. evolved PCA) calculated from the empirical evolved trait values from all independent populations in a *Chlamydomonas* long-term evolution study [3]. The starting coordinate for the initial population (tan circle in Fig. 2b) was chosen based on the trait values of a single *Chlamydomonas* ancestral genotype projected onto the evolved trait-scape. The evolutionary end-point, or the high fitness area (red circle in Fig. 2b), was chosen based on the trait values of that same *Chlamydomonas* genotype following $CO_2$ adaptation in the evolved trait-scape. Below we outline the steps for each generation in a single model run.

In the default model simulations (referred to as 90/10), 90% of individuals were randomly chosen to experience a random change in a trait value (while maintaining all existing trait-correlations) while 10% experienced both a trait and trait-correlation change. Each change was drawn from a Gaussian distribution (mean = 0 and standard deviation = 0.05) such that small changes were common and large changes were rare. For each individual, the randomly chosen trait change was added to the existing trait value. Following this initial trait change, the remaining 3 trait values were updated using the trait correlations for that individual in that time step. For example, if trait 1 was initially changed, then traits 2, 3, and 4 would subsequently be updated by multiplying the new trait 1 value by the three trait correlations (1v2, 1v3, and 1v4) for that individual.

The remaining 10% of the population experienced both a trait and a trait correlation change. For each individual, one of the six trait correlations was randomly selected to change. Similar to the trait change, a random value was drawn from a Gaussian distribution with a mean of 0 and standard deviation of 0.05 and added to the existing correlation value. Next, one of the two traits associated with that correlation was randomly chosen and a trait change was selected in the same manner as above. Next, the second trait tied to the correlation was updated using the new correlation and trait value (the other 2 trait values were not updated in this generation). Following these changes, each individual was projected back onto the evolved trait-scape (i.e., evolved PCA) using the evolved factor loadings. This generated movement of each individual within the trait-scape.

Selection was then imposed by first calculating the Euclidean distance between the location of each individual in the trait-scape and the high fitness area (evolutionary end coordinate). Fitness for each individual was then calculated as a function of the euclidian distance ($z$) to the high fitness area as:

$$w(z) = e^{(-z^2)/2} \quad \text{Eq. 1}$$

Finally, individuals were randomly sampled with replacement weighted by these new fitness values to persist to the next generation. As a result, individuals with low fitness were more likely to go extinct (not persist to the next generation) while individuals with high fitness were more likely to persist (or even increase their representation) in the next generation.

**Supplementary Information 4:** ***Integrating the TRACE approach into an ODE framework***

Biogeochemical models commonly assume fixed trade-offs between traits, for example, size and metabolic rate [4-6]. Ultimately, these trade-offs drive the cycling of nutrients in the model and the resulting community diversity. For example, small phytoplankton typically have maximum growth rates compared to large cells [7,8]. However, there are many traits for which there is only a weak (or no) relationship with size. For example, while maximum uptake rate is highly correlated with cell size, nutrient affinity only shows a very weak relationship to cell size[9,10]. Similarly, while Qmin shows a strong relationship with cell size, there is a very weak relationship between cellular C:N or Qmax:Qmin and cell size [11]. A weak relationship is also observed between cell size and light harvesting traits [12,13], and several experimental evolution studies have shown that cells can evolve light harvesting traits to increase photosynthetic performance thereby changing the relationship between size and light harvesting traits [14-17].

Overall there is increasing evidence that most trait-trait relationships can be modified under selective pressure, though some more than others, and that trends based on interspecific observations may not provide much constraint on intraspecific change. Furthermore, even small changes in trait correlations (e.g., between maximum growth rate and cell size) can result in significant changes to ecosystem structure particularly in fluctuating environments [18]. Therefore, to understand how plankton will evolve we need a new modeling framework that accounts for evolutionary change, including how the relationship between traits may evolve. TRACE could provide such a framework when coupled to an ecosystem model. While a full coupling of TRACE and an ecosystem model are beyond the scope of this paper, here we provide some sensitivity studies to demonstrate the significance of such a coupling.

*Ecosystem model:* Here we use a standard trait-based phytoplankton model modified from the DARWIN model [6]. The model simulates the growth of phytoplankton considering light and nutrient limitation and allows for phytoplankton plasticity by allowing their nitrogen and chlorophyll to carbon quotas to be emergent properties. As in Ward and Follows (2016) [6], photosynthesis is described as an exponential function of irradiance, chlorophyll to carbon ratio, and alpha (initial slope of the photosynthesis irradiance curve). The uptake of inorganic nitrogen is defined using a Michaelis-Menten equation and the nutrient status of the cell, so that the uptake of nutrients is reduced as the quota becomes full. In our model, $\mu_{max}$ ultimately defines the maximum inorganic nutrient uptake rate ($V_{max}$, gN $gC^{-1}$ $day^{-1}$), the maximum photosynthesis rate ($P_{max}$, gC $gC^{-1}$ $day^{-1}$), and the respiration rate ($R_C$, gC $gC^{-1}$ $day^{-1}$), following Flynn and Mitra (2009) [19]:

$$V_{max} = \mu_{max}\, NC_{max}$$

$$P_{max} = \mu_{max} + \mu_{max}\, basres + \mu_{max}\, NC_{max}\, biosyn\, \gamma_N$$

$$R_C = \mu_{max}\, basres + biosyn\, N_{up}$$

where $NC_{max}$ is the maximum nitrogen to carbon quota, *basres* is the relative basal respiration cost, *biosyn* is the cost associated to nutrient uptake (gN $gC^{-1}$), $\gamma_N$ is the nutrient limitation factor and $N_{up}$ is the actual nutrient uptake rate (gN $gC^{-1}$ $day^{-1}$). We also included a quadratic mortality term to represent non-resolved losses (e.g. predation).

*Model simulations:* We used the ecosystem model to illustrate how evolutionary adaptation of the relationships between traits (trait correlations) has the potential to change the growth dynamics and the competitive abilities of phytoplankton. We ran two versions of the model: ancestral correlations in which both $\mu_{max}$ and α are inversely related to cell size based on interspecific allometries from the literature [13,20]; and evolved correlations in which large cells have evolved higher $\mu_{max}$ and $\alpha$ at the cost of higher respiration rates (i.e. a change in the trait-correlations for $\mu_{max}$, $\alpha$, and respiration). These evolved correlations were chosen based on an experimental evolution study which showed that large cells can achieve higher photosynthetic performance and growth rates than small cells but that the higher demand for chlorophyll synthesis and photosynthetic efficiency in general resulted also in higher respiration rates for the large cells [16]. In addition, the large cells showed lower oxygen production per chlorophyll molecule (as expected by the package effect) [16]. The parameter values for the two versions of the model are given in Supplementary Table 2.

The model was run with two phytoplankton groups, small (100 $\mu m^3$) and large (1000 $\mu m^3$) cells, competing for resources in a chemostat with a constant dilution rate (0.05 $day^{-1}$). A 20x20 matrix of environmental conditions were tested with 400 combinations of irradiance (PFD: 50–600 $\mu$mol photon $m^{-2}$ $day^{-1}$) and dissolved inorganic nitrogen (DIN: 1–100 $\mu$g N $L^{-1}$) values. All simulations were run to steady-state and the growth dynamics of the two different size classes were analyzed.

*Model results:* When ancestral correlations were applied, small cells achieved higher biomass levels than large cells across all levels of light and nutrients (Supplementary Fig. 6a). This is due to the ancestral allometric rules (based on interspecific comparisons from the literature) which imply that small cells have higher light affinity and maximum growth rates than large cells. However, when the large cells were allowed to evolve the relationships between their traits, we see the relative biomass levels for the two groups shift (Supplementary Fig. 6b). Specifically, by evolving higher $\mu_{max}$ and $\alpha$, large cells now have higher growth capacity under non-limiting conditions compared to small cells. However, as $\mu_{max}$ and $\alpha$ are positively correlated with respiration costs for the large cells, small cells can achieve higher biomass compared to large cells under resource limitation. By looking at the biomass ratio between small and large cells (Supplementary Fig. 7), we see that large cells are the winner (values < 1) when nutrients are not limiting (though this advantage decreases as light becomes severely limiting) and small cells are the winner (values > 1) when nutrients are limiting growth.

This simple example shows the potential of evolving trade-offs and trait correlations to change the growth environment of phytoplankton and their competitive abilities in models. Ultimately, the quantitative framework of TRACE, informed by trade-off data from evolution experiments, will allow us to better constrain the evolution of multiple trait correlations in plankton models.

**Supplementary Table 1** Parameter values used for main text model simulations.

| Parameter | Description | Value |
|---|---|---|
| ***N*** | Size of population | 1000 |
| ***t*** | Number of generations | 2000 |
| ***tgrad*** | Standard deviation of trait change | 0.05 |
| ***cgrad*** | Standard deviation of correlation change | 0.05 |
| $\mathbf{N_{trait}}$ | Number of trait changes | [900, 500, 100] |
| $\mathbf{N_{corr}}$ | Number of correlation changes | [900, 500, 100] |
| $\mathbf{N_{runs}}$ | Number of replicate runs | 100 |

**Supplementary Table 2** Parameter values determining the relationships between traits for small (100 μm$^3$) and large (1000 μm$^3$) cells.

| Scenario | Cell size | $\mu_{max}$ (gC gC$^{-1}$ day$^{-1}$) | α (gC gChl$^{-1}$ × m$^2$ μmol photon$^{-1}$) | biosyn (gN gC$^{-1}$) |
|---|---|---|---|---|
| Ancestral correlations | Small | 1.2 | $1 \times 10^{-5}$ | 3.2 |
| | Large | 0.7 | $4 \times 10^{-6}$ | 3.2 |
| Evolved correlations | Small | 1.2 | $1 \times 10^{-5}$ | 3.2 |
| | Large | 1.4 | $8 \times 10^{-6}$ | 6.4 |

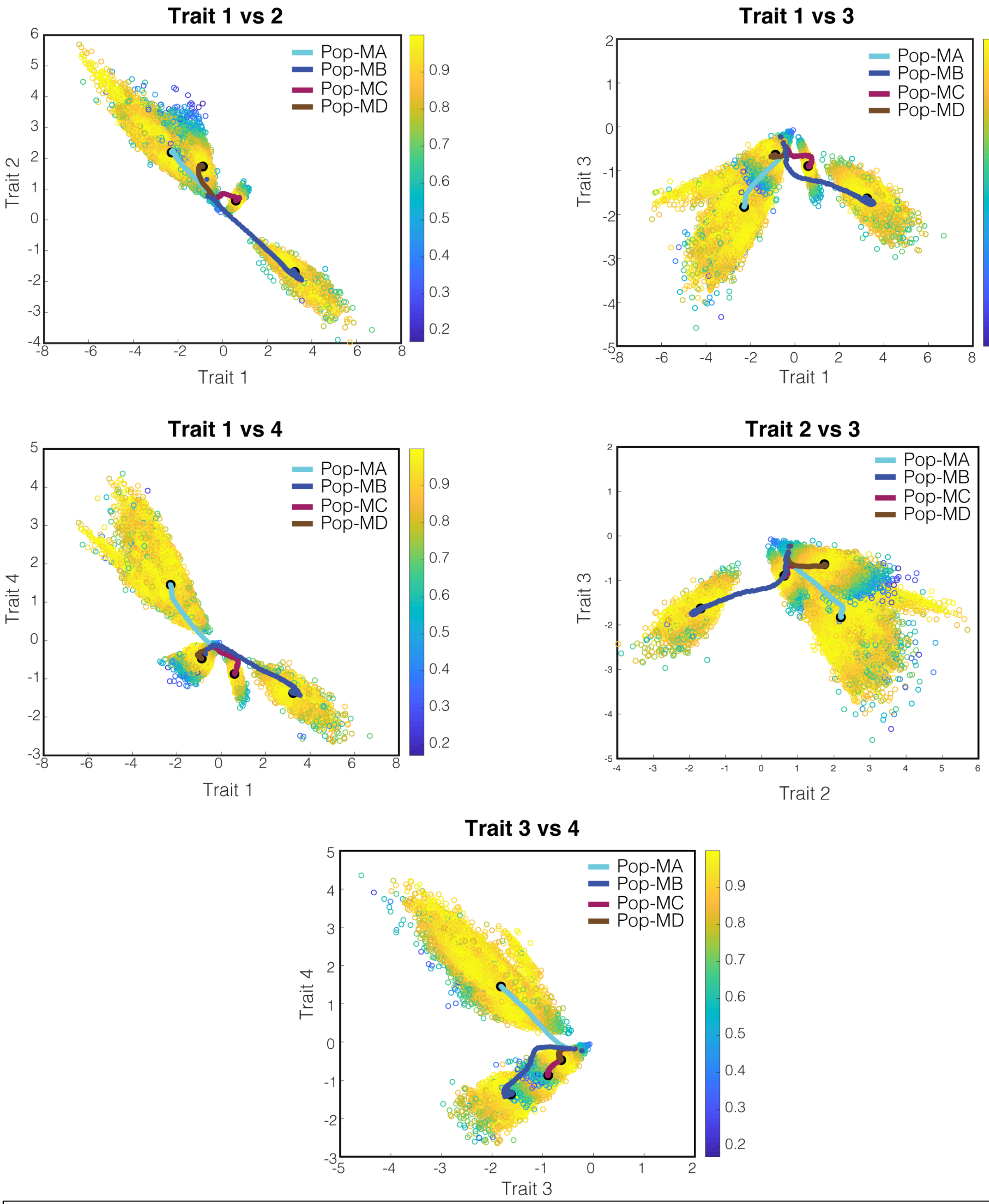


**Supplementary Fig. 1 Trait vs. trait plots for mixed mode simulations** a) Trait vs. trait plots denoting the 4 distinct population types (i.e. cryptic phenotypes) that emerged from 100 replicate model runs in mixed mode (i.e., no bias). Each hollow point represents the final trait values of a given individual in the last generation (2000$^{th}$) colored by fitness. Colored lines represent the average trait values at each generation for each population with the black point denoting the final generation.

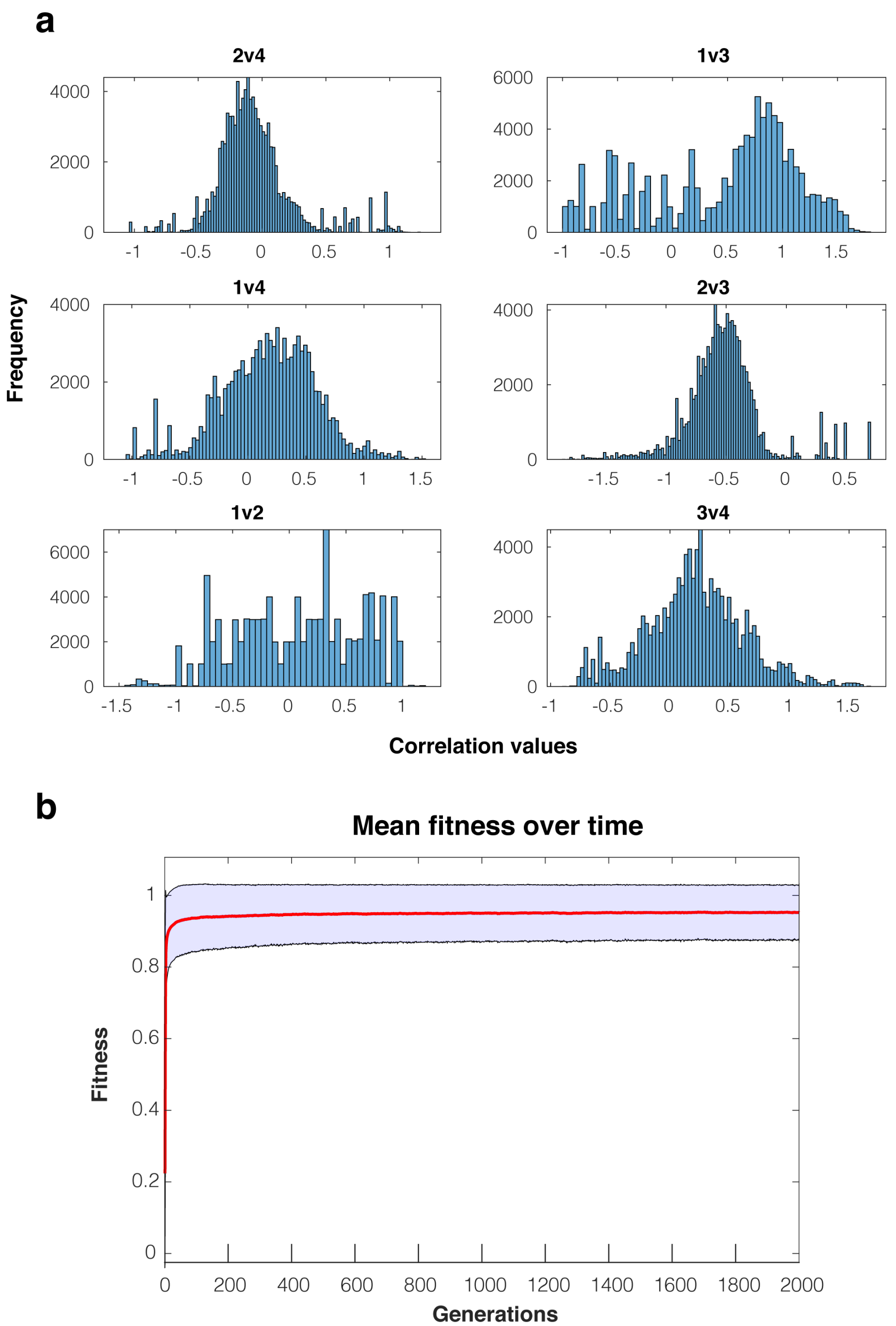


**Supplementary Fig. 2 Trait correlational distributions from model runs that changed traits independently of trait correlations.** a) Shown are trait correlational distributions per correlation. At each generation, 1 trait per individual was randomly changed but no other traits were updated using the trait correlations resulting in one phenotypic solution. b) Shown is the mean fitness of all individuals over time across all replicate runs (1000 individuals x 100 replicate runs). The red line indicates the mean while the light blue area denotes the smoothed standard deviation.

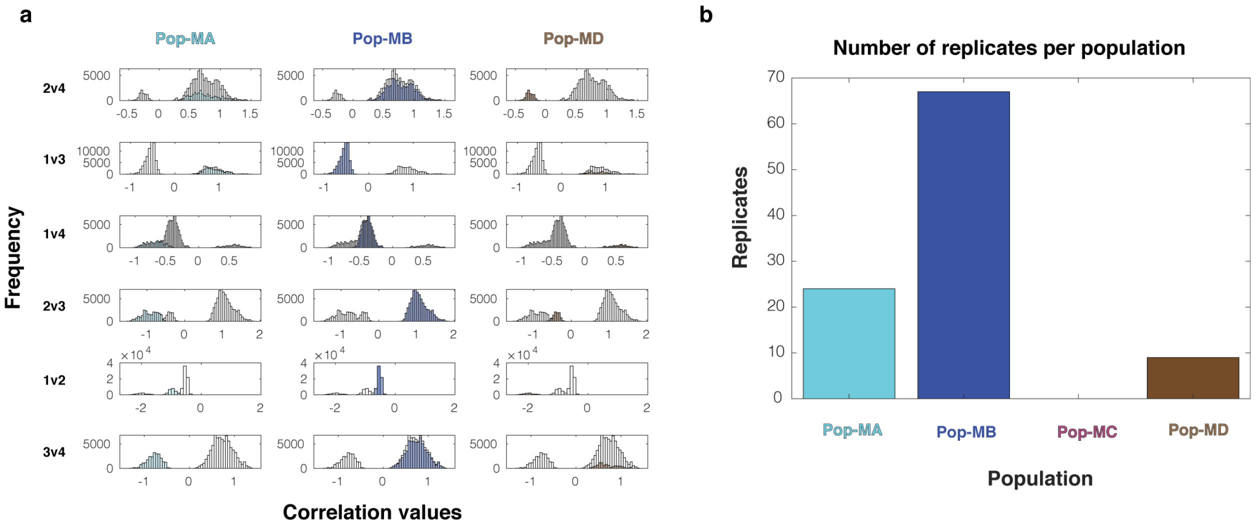


**Supplementary Fig. 3 Impact of starting location on emergent population types (i.e. cryptic phenotypes) in mixed-mode model simulations.** a) Shown are emergent population types resulting from mixed-mode model simulations as in the main text except starting from a different, equidistant ancestral phenotype in PCA space. Each row displays one of the six possible trait correlations (2v4, 1v3, 1v4, 2v3, 1v2, and 3v4) with the distribution of the emergent trait correlation values for all individuals in all replicate runs (N=100,000) shown in grey. Highlighted in color in each subplot are the trait correlation values for the individuals belonging to each of the emergent population type (columns). Each population type has a clearly defined set of trait correlation values. b) Shown is a bar plot denoting the number of replicates within each population type (out of 100 model runs). When starting from a different ancestral start point, Pop-MC did not emerge in any replicates.

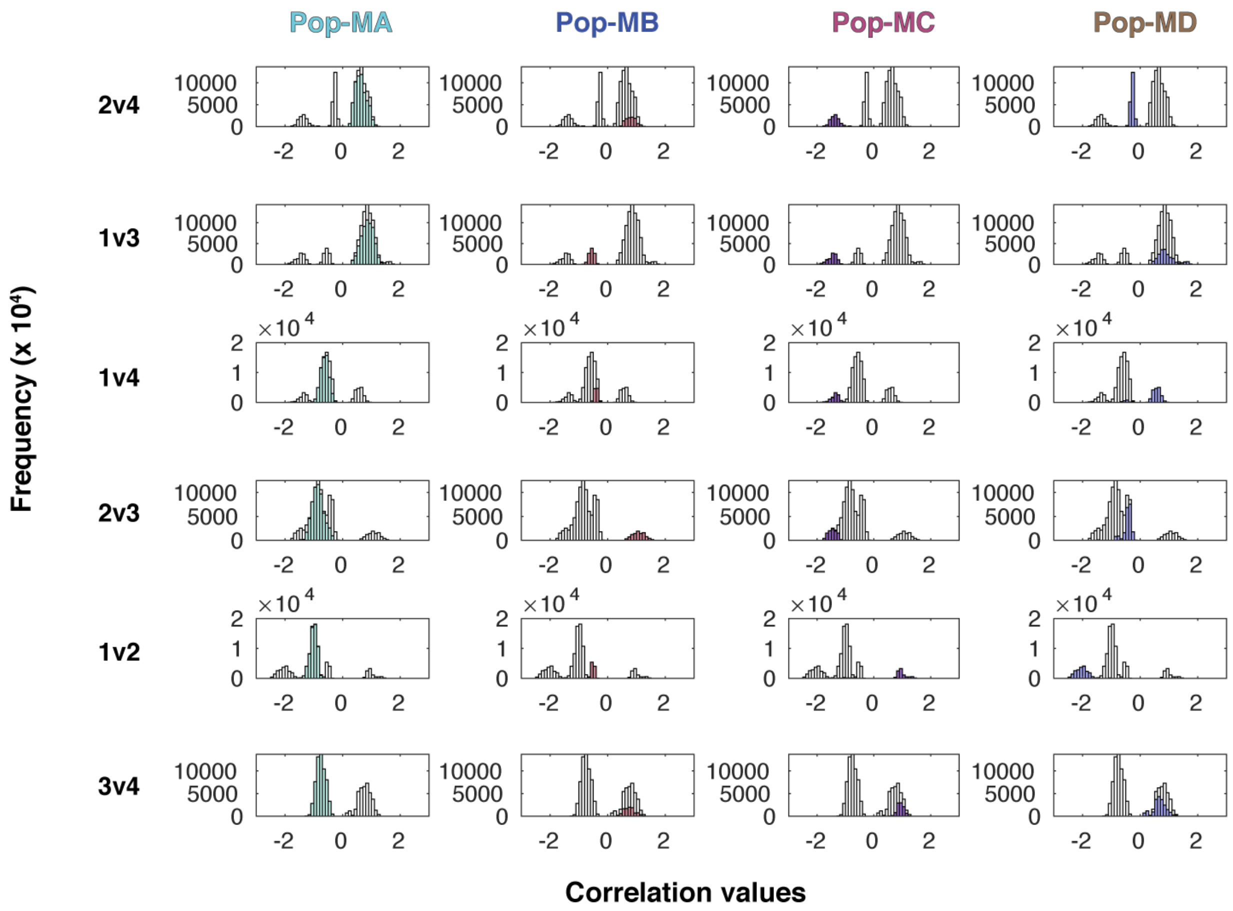


**Supplementary Fig. 4 Four distinct, emergent population types (i.e. cryptic phenotypes) from model runs seeded with 4 different subpopulations and no bias** These model runs were identical to "Mixed mode" in the main text (Fig. 4) except that instead of starting with a clonal phenotype (same starting trait values), we initialized 4 different subpopulations (i.e. 4 different starting locations). Each row displays one of the six possible trait correlations (2v4, 1v3, 1v4, 2v3, 1v2, and 3v4) with the distribution of the emergent trait correlation values for all individuals in all replicate runs (N=100,000) shown in grey. Highlighted in color in each subplot are the trait correlation values for the individuals belonging to each of the emergent population type (columns). Each population type has a clearly defined set of trait correlation values. Here, the 4 same populations emerged as in Fig. 4 of the main text.

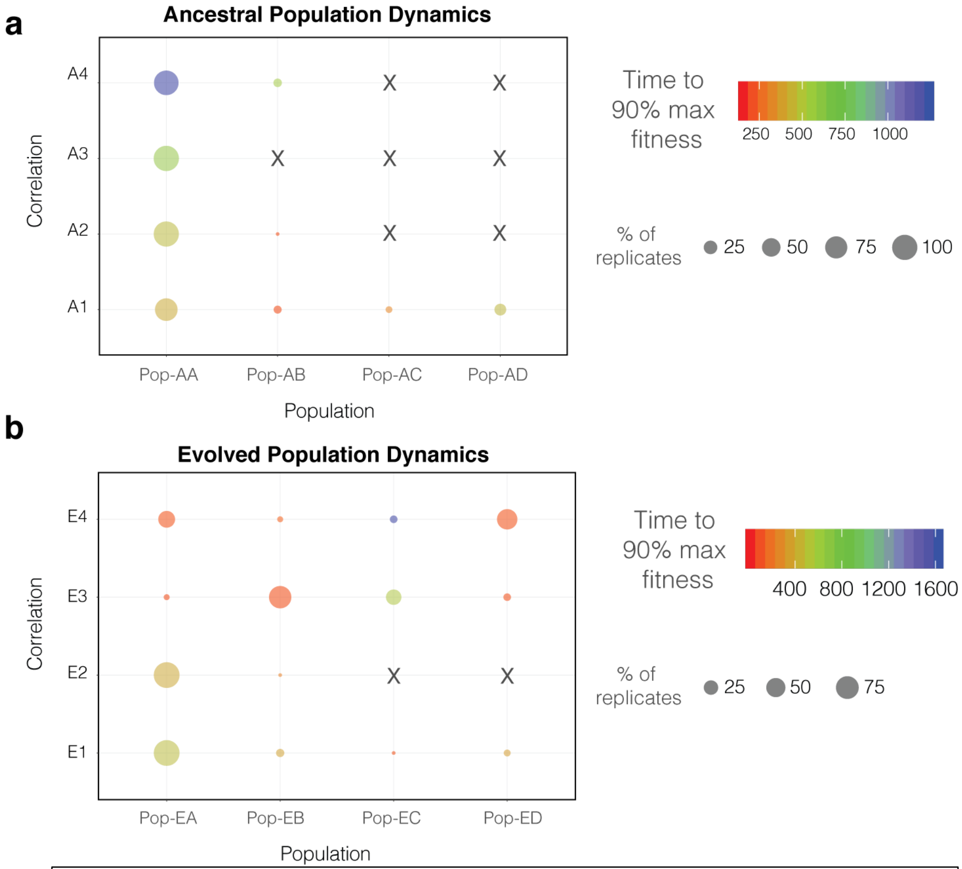


**Supplementary Fig. 5 Dynamics of emergent population types across different historical bias** a) Bubble plot showing emergent population types as a function of adding empirical ancestral correlations (ancestral bias). Bubble size denotes the number of replicates (out of 100) within a specific population type while bubble color represents each population's rate of adaptation (color of circles). b) Same plot as in a) except with adding empirical evolved correlations (evolved bias).

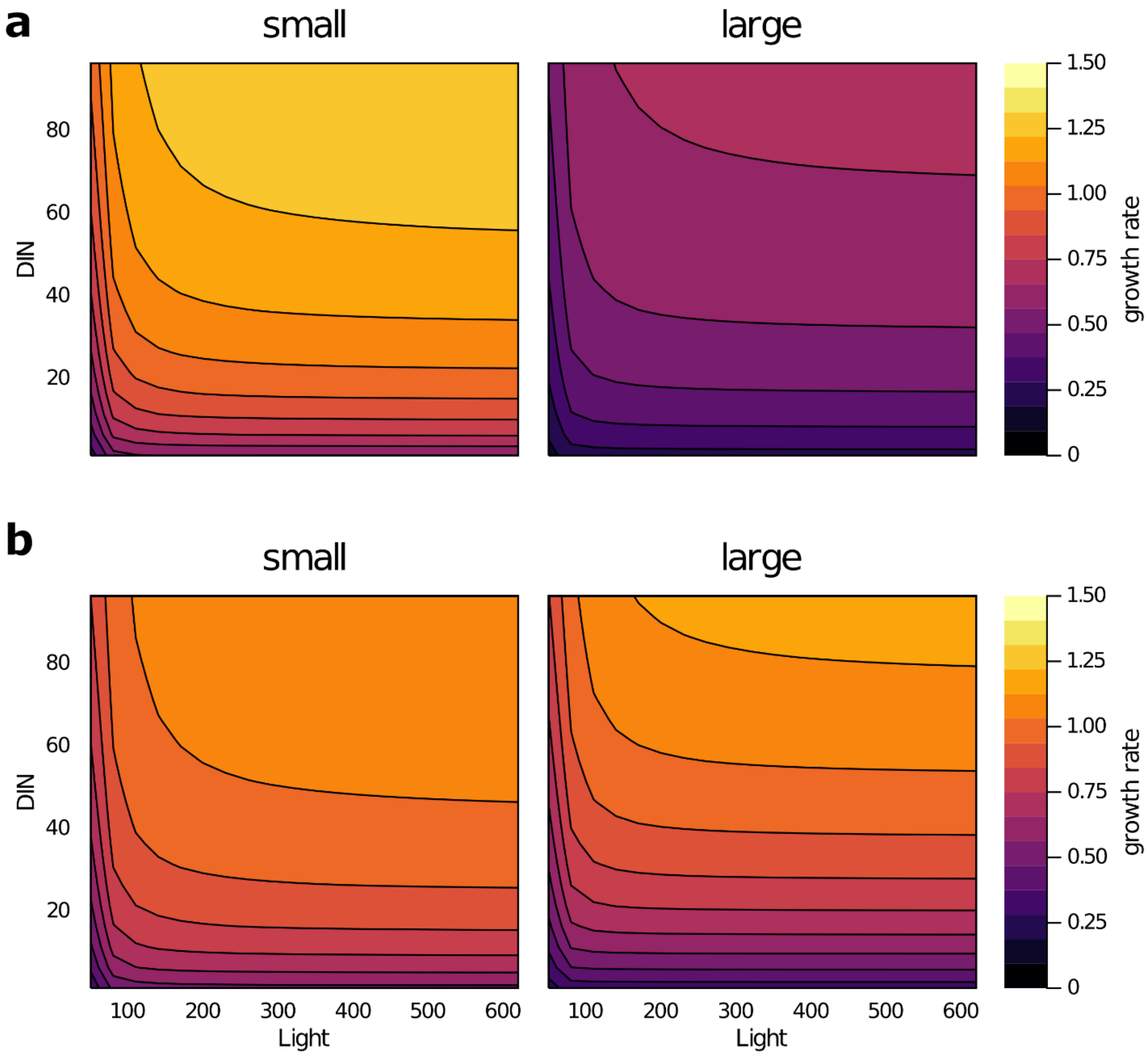


**Supplementary Fig. 6 Growth dynamics of two phytoplankton of different sizes (small = 100 µm3 and large = 1000 µm3) competing for the same resources at steady state.** Results are shown for the scenarios of ancestral correlations (a) and evolved correlations (b). Contour plots were obtained for a matrix of 20x20 values of DIN: dissolved inorganic nitrogen (µg N L-1) and light (µmol photon m-2 day-1). Growth rate (gC gC-1 day-1) is given by the color bars.

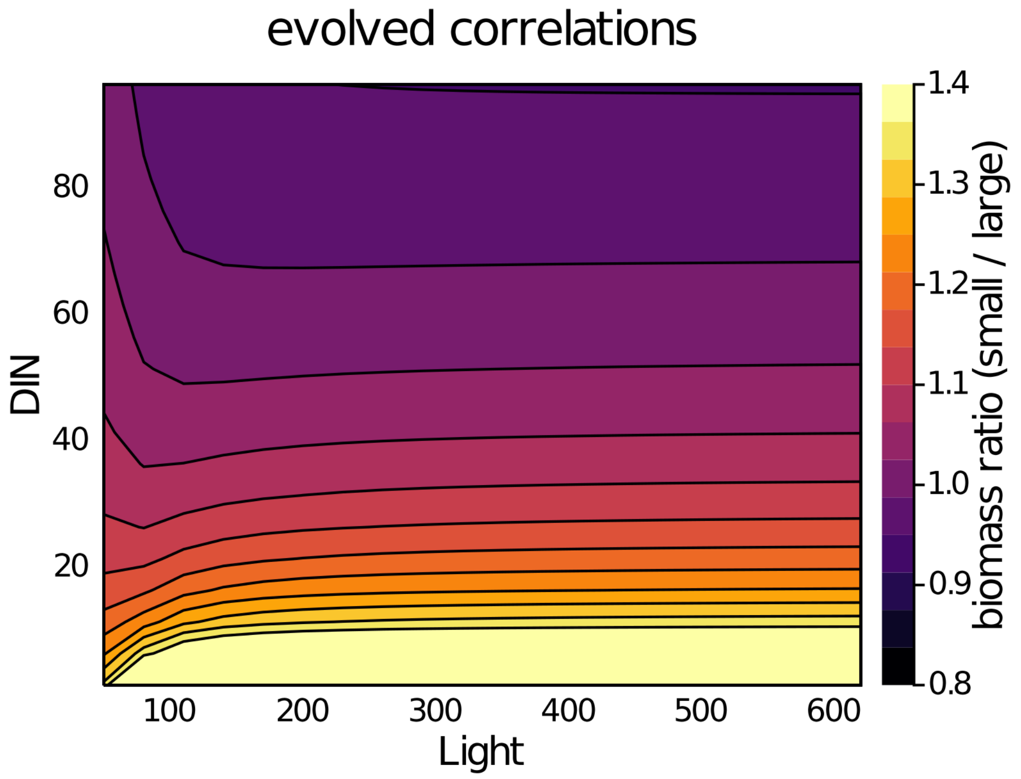


**Supplementary Fig. 7 Relative success of two phytoplankton of different sizes (small = 100 μm$^3$ and large = 1000 μm$^3$) competing for the same resources at steady state**. The biomass ratio between the small and large phytoplankton populations is given by the color bar. Results are shown for the scenario of evolved correlations. While small cells were always the winners in the simulations with ancestral correlations, the large cells were able to gain an advantage in some environments by evolving their trait correlations. Contour plots were obtained for a matrix of 20x20 values of DIN: dissolved inorganic nitrogen (μg N L-1) and light (μmol photon m-2 day-1).

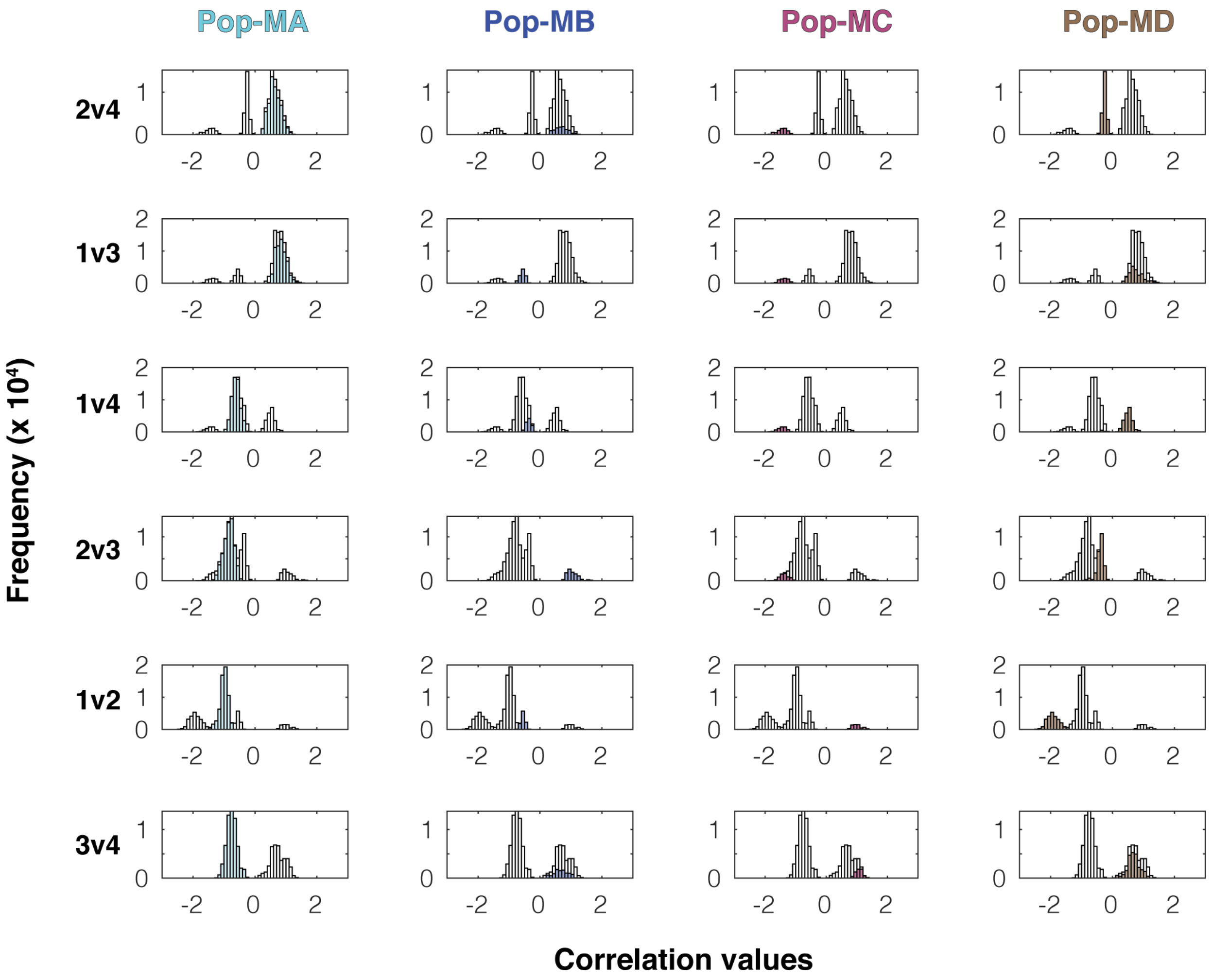


**Supplementary Fig. 8 Four distinct, emergent population types (i.e. cryptic phenotypes) from model runs seeded with no bias** These model runs were identical to "Mixed mode" in the main text (Fig. 4) except the order of updating traits and correlations was reversed. Each row displays one of the six possible trait correlations (2v4, 1v3, 1v4, 2v3, 1v2, and 3v4) with the distribution of the emergent trait correlation values for all individuals in all replicate runs (N=100,000) shown in grey (y-axis, note scale of $10^4$). Highlighted in color in each subplot are the trait correlation values for the individuals belonging to each of the emergent population types (columns). Each population type has a clearly defined set of trait correlation values. Here, the 4 same populations emerged as in Fig. 4 of the main text.

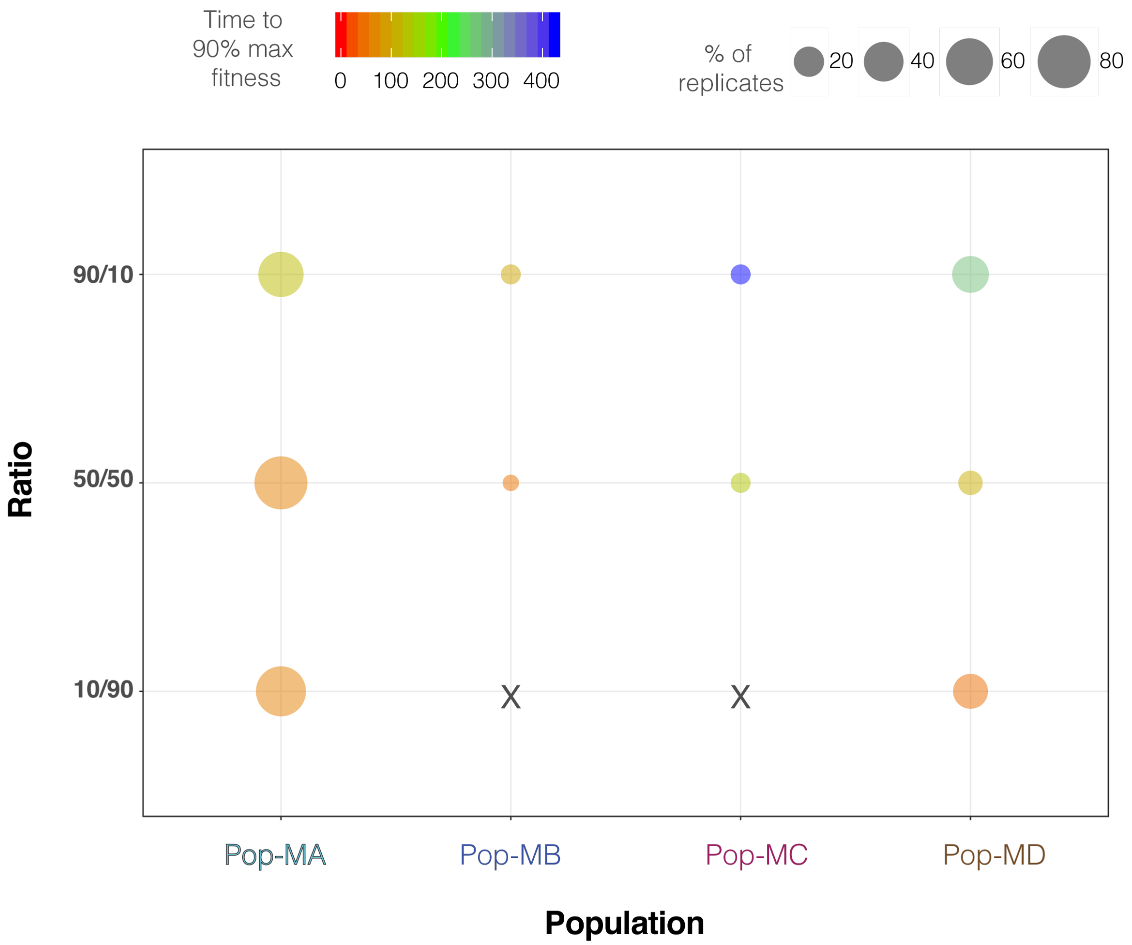


**Supplementary Fig. 9 Population dynamics of emergent population types across different ratios of trait and trait correlation changes.** The bubble plot shows emergent population types as a function of varying the ratio of trait and trait correlation changes for mixed-mode simulations. 90/10 denotes 90% of individuals in a population experiencing a trait change and 10% both a trait and correlation change per generation. Similarly, 50/50 denotes 50% experiencing a trait change and 50% both a trait and correlation change. Finally, 10/90 denotes 10% experiencing a trait change and 90% both a trait and correlation change. Bubble size denotes the number of replicates (out of 100) within a specific population while bubble color represents each population's rate of adaptation (color of circles), or time to reach 90% of its maximum fitness.
